# A portable, easy-to-use paper-based biosensor for rapid in-field detection of fecal contamination on fresh produce farms

**DOI:** 10.1101/2023.10.04.560915

**Authors:** Jiangshan Wang, Simerdeep Kaur, Ashley Kayabasi, Mohsen Ranjbaran, Ishaan Rath, Ilan Benschikovski, Bibek Raut, Kyungyeon Ra, Nafisa Rafiq, Mohit S. Verma

**Author notes:** Corresponding author’s.

## Abstract

Laboratory-based nucleic acid amplification tests (NAATs) are highly sensitive and specific, but they require the transportation of samples to centralized testing facilities and have long turnaround times. During the Coronavirus Disease 2019 (COVID-19) pandemic, substantial advancement has been achieved with the development of paper-based point-of-care (POC) NAATs, offering features such as low cost, being easy to use, and providing rapid sample-to-answer times. Although most of the POC NAATs innovations target clinical settings, we have developed a portable, paper-based loop-mediated isothermal amplification (LAMP) testing platform for on-farm applications, capable of detecting *Bacteroidales* as a fecal contamination biomarker. Our integrated platform includes a drop generator, a heating and imaging unit, and paper-based biosensors, providing sensitive results (limit of detection 3 copies of *Bacteroidales* per cm^2^) within an hour of sample collection. We evaluated this integrated platform on a commercial lettuce farm with a concordance of 100% when compared to lab-based tests. Our integrated paper-based LAMP testing platform holds great promise as a reliable and convenient tool for on-site NAATs. We expect that this innovation will encourage the fresh produce industry to adopt NAATs as a complementary tool for decision-making in growing and harvesting. We also hope that our work can stimulate further research in the development of on-farm diagnostic tools for other agricultural applications, leading to improved food safety and technology innovation.

## 1. Introduction

In recent years, lab-based nucleic acid amplification tests (NAATs), such as quantitative polymerase chain reaction (qPCR), have been used to detect foodborne pathogens (Dinu & Bach, 2013; Elizaquível et al., 2012). Lab-based NAATs necessitate the transportation of obtained field samples to centralized testing labs, where the samples are tested by highly experienced technicians using qPCR assays on expensive thermocyclers. However, lab-based NAATs are not economical due to the additional high expenses (e.g., cold-chain overnight shipping, lab-testing service fee) and time delay (usually 24 – 48 hours), which may impair the freshness and marketability of products (Paniel & Noguer, 2019). Timely response from testing in the fresh produce industry is important because fresh produce has a limited shelf life. Rapid testing allows for quick identification and remediation of any potential food safety issues, reducing the risk of spoilage and food waste. There is a need for rapid, simple-to-use, inexpensive, highly sensitive in-situ testing. With minimal training requirements, non-specialist users can use these tests in the field with simple equipment to determine site-specific risks and assist fresh produce growers in their decision-making process regarding the microbial safety of fresh produce (Arora et al., 2011).

Furthermore, in the majority of cases, the concentration of the enteric pathogens is relatively low and it is not realistic for nucleic acid testing alone to meet the requirement of sensitivity for foodborne pathogens detection. Limit of detection (LoD) for qPCR at 95% confidence interval is 0.15 copies/µL (3 copies of target in a 20 µL reaction) (Forootan et al., 2017); the required sensitivity for foodborne pathogens detection assays is 4.4×10^-6^ copies/µL (1 copy diluted in 225 mL of broth, without enrichment) (FDA, 2021; Ferone et al., 2020). As a result, the qPCR assay is used with culture-based enrichment methods, which can run into the problem of pathogens entering a viable-but-nonculturable (VBNC) state (FDA, 2021). Aside from determining the presence or absence of pathogens, an alternative is to quantify the abundance of fecal indicator bacteria (FIB)—microorganisms selected as indicators of fecal contamination (Brauwere et al., 2014; Wang, Ranjbaran, Ault, et al., 2023). FIB such as *Escherichia coli*, *Enterococcus faecalis*, and *Bacteroidales*, have been used to assess possible fecal contamination in fresh produce (Denis et al., 2016; Drozd et al., 2013; Harris et al., 2017; Ordaz et al., 2019; Wang, Ranjbaran, & Verma, 2023). Compared to other FIB, *Bacteroidales* is preferable due to the following four features: i) high prevalence in feces (constituting 30%–40% of total fecal bacteria, 10^9^ to 10^11^ CFU/g), ii) low natural abundance from non-fecal sources, iii) obligate anaerobicity (preventing their growth and multiplication in the ambient environment), and iv) high host specificity (host-specific markers on the 16S rRNA gene can be used for microbial source tracking) (Mascorro et al., 2018; Ordaz et al., 2019). As a result, *Bacteroidales* serve as a valuable—and potentially quantitative—marker.

During the COVID-19 pandemic, unprecedented research effort has resulted in the significant development of paper-based point-of-care (POC) NAATs (Sritong et al., 2023; Wang et al., 2022). These analytical devices provide several benefits over conventional lab tests including lower costs, simple fabrication processes, short sample-to-answer time, and user-friendliness (Sritong et al., 2023; Wang et al., 2022). By preloading the paper-based biosensor with amplification reagents and implementing a smart design, a multistep amplification process can be completed on one chip, improving user-friendliness for non-specialist users. Isothermal amplification methods, such as loop-mediated isothermal amplification (LAMP), enable single-temperature operations and can be performed with simple equipment such as a water bath or incubator (Davidson et al., 2021; De Felice et al., 2022; Dinh & Lee, 2022; Kang et al., 2022; Mohan et al., 2021; Notomi et al., 2000; Pascual-Garrigos et al., 2021; Wang et al., 2021, 2022; Wang, Ranjbaran, Ault, et al., 2023).

While none of the previous works have focused on enabling LAMP testing for on-farm applications, our group made several efforts in this direction (Pascual-Garrigos et al., 2021; Wang, Ranjbaran, Ault, et al., 2023). We have shown that LAMP has the potential for on-farm NAAT and have established paper-based assays as a method that can partly decrease the number of steps for performing *in-situ* tests (Davidson et al., 2021; Pascual-Garrigos et al., 2021; Wang et al., 2021; Wang, Ranjbaran, Ault, et al., 2023). We also developed and tested a micro-drop generator as another effort to enhance user-friendliness (Ranjbaran et al., 2023). By pressing a single button, the micro-drop generator delivers a precise volume of sample to the paper-based biosensor, eliminating the need for commercial micro-pipettors.

In the present study, we utilized all the previous advancements to implement paper-based LAMP testing in fresh produce farms. We developed a fully-integrated LAMP testing platform which included components such as heating, imaging, fluid delivery, and paper-based LAMP assay, and deployed it on a commercial lettuce farm. The unit, operating at the back of a car, was powered by a portable power station and detected *Bacteroidales* as the biomarkers of fecal contamination. Our platform enabled *in-situ* identification of fecal contamination within 60 min of sampling. To our knowledge, this study is the first demonstration of implementing a portable paper-based LAMP testing platform in a fresh produce farm. It serves as an enabler for establishing future NAATs as part of standard growing and harvesting practices during fresh produce production.

## 2. Materials and methods

### 2.1 Design and fabrication of Field-Applicable Rapid Microbial Loop-mediated isothermal Amplification Platform (FARM-LAMP)

All designs were prepared using SolidWorks software (SolidWorks, MA). The heating unit consists of a 3D printed cavity, made out of Dental LT Clear V2 resin (Formlabs, RS-F2-DLCL-02) for the first FARM-LAMP device and Biomed Clear V1 resin (Formlabs, RS-F2-BMCL-01) for the second FARM-LAMP device, with a 3 mm transparent acrylic sheet (Amazon, B099J2XVRW) attached at its bottom (Figure 1A). The platform holding the imaging components was printed with Grey Pro V1 resin (Formlabs, RS-F2-PRGR-01) for the first FARM-LAMP device and Raise 3D Premium PLA filament in art white (Raise3D, CA) for the second FARM-LAMP device. The first and second FARM-LAMP devices have very similar designs, except the second device has a small cooling fan (Amazon, B06XQDMMJ5) installed on the bottom of the platform. Different printing materials were used due to availability at the time of manufacturing. All resin-based 3D printing was performed using a Form 3B stereolithography 3D printer (Formlabs, MA) while filament-based 3D printing was performed with a Raise3D Pro2 Plus 3D printer (Raise3D, CA). The transparent window under the heating cavity allowed for coupling the heating unit with an imaging unit to track the color changes in the reaction pads. Each FARM-LAMP device used two 80W, 120 V hot rod Dernord heating elements (Amazon, B08LK9HCWW) to heat the water as well as two 12 V submersible mini water pumps (Amazon, B08RWP6GJF) to circulate water in the tank and improve the temperature uniformity. The water temperature was monitored using a waterproof digital temperature sensor (Gikfun, DS18B20), controlled by a PID control algorithm, and ran on a Raspberry Pi 4B (Amazon, B07TD42S27) mini-computer. The FARM-LAMP device provided fast operation, reaching 65 °C in under 20 minutes. A high resolution autofocus camera with 16 megapixel imaging resolution (Amazon, B09STL7S88) in the imaging unit captured a shot of the paper pads every minute.

**Figure 1:**
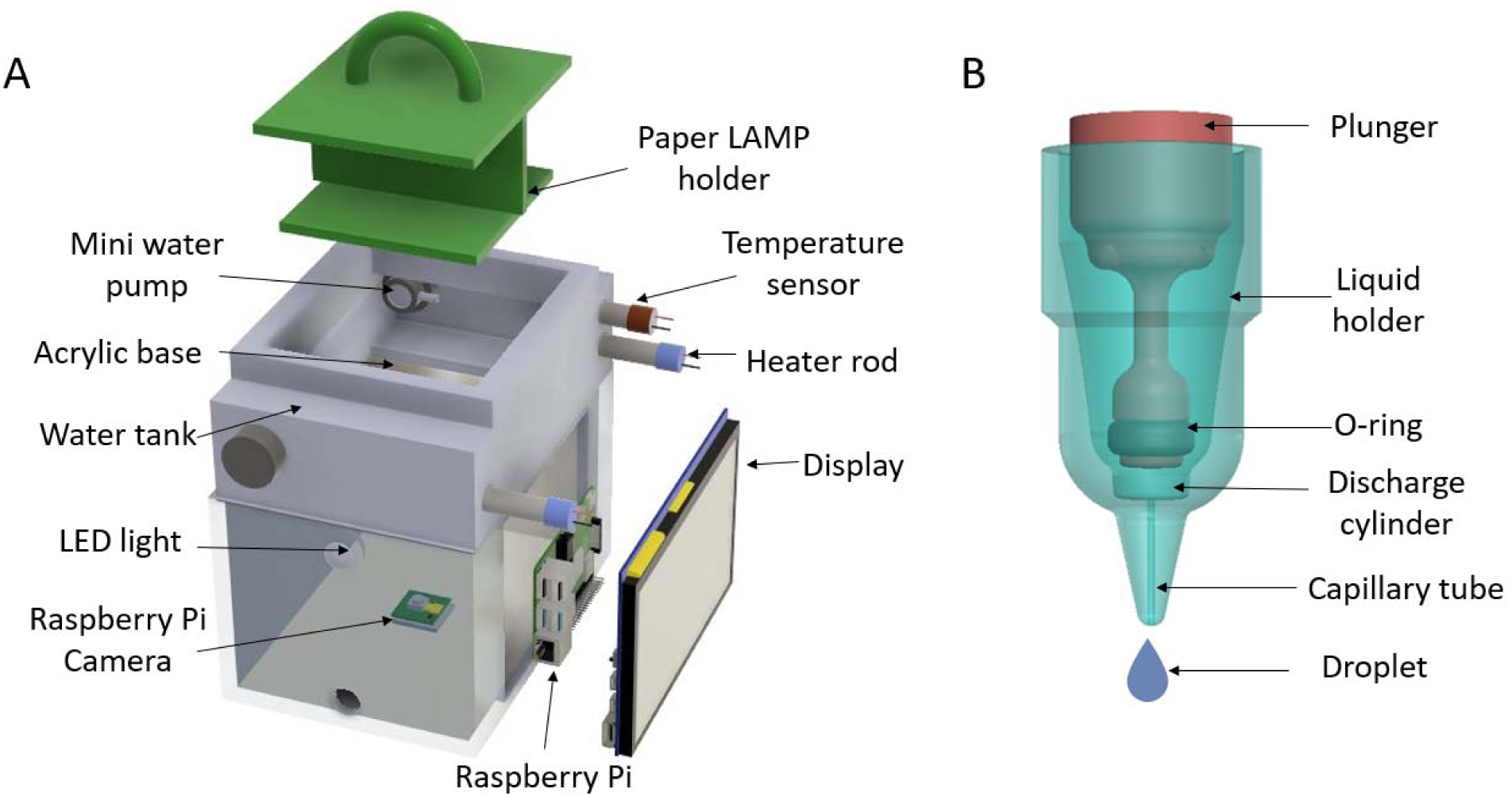
Design of the heater (A) and drop generator (B).

### 2.2 Fabrication of drop generators

To measure and create precise volumes of the microbial samples in the field, we designed a drop generator with a capacity of 27 µL of drop volume. A comprehensive description of the drop generators and their efficacy was provided elsewhere (Ranjbaran et al., 2023). Briefly, the generator consisted of a liquid holder, a plunger, and two O-rings (Helipal, Airy-Acc-Oring-2.5×6 mm) (Figure 1B). The liquid holders and plungers were 3D printed using High Temp V2 resin (Formlabs, RS-F2-HTAM-02). After printing, the parts were washed with isopropyl alcohol (IPA) for 30 minutes and cured under UV light at 55 °C for 30 minutes. The tip of the liquid holders was wrapped in Parafilm^TM^ wrapping film prior to surface treatment (Fisher Scientific, S37440). The liquid holders were surface-treated with oxygen plasma for 2 minutes at 0.2 Torr using a plasma generator (Plasma Etch, Inc., PE-25) and polyethylene glycol (PEG) 400 (Fisher Scientific, P167-1) for 24 hours. Following surface treatment, all components were washed with IPA, then with ultrapure water (PURELAB flex, ELGA), dried with an air gun, and stored in separate resealable polypropylene bags (Uline, S-17954) for future use.

### 2.3 Fabrication and preparation of microfluidic paper-based devices (µPADs)

µPADs were used for running LAMP assays in the field (Wang et al., 2021). Each device consisted of two paper pads of 5 mm × 6 mm chromatography paper (Ahlstrom-Munksjo, Grade 222) attached on a double-sided adhesive (Adhesives Research, 90178). The paper pads were separated by 5 mm polystyrene spacers (Tekra, 40047020). After fabrication, the µPADs were stored in resealable polypropylene bags (Uline, S-17954) for future use.

### 2.4 Preparation of LAMP reaction mix and pads

The LAMP reaction mix reported by Wang *et al*. was used here (Wang et al., 2021). To prepare 1000 µL 2x LAMP mix, the solution consisted of 100 µL KCl (1000 mM; Sigma-Aldrich, P9541), 160 µL MgSO_4_ (100 mM; Sigma-Aldrich, M2773), 280 µL deoxynucleotide triphosphate (dNTP) (10 mM; Fisher Scientific, FERR0182), 2.8 µL deoxyuridine triphosphate (dUTP) (100 mM; Fisher Scientific, FERR0133), 0.4 µL Antarctic Thermolabile UDG (1 U/µL; New England Biolabs, M0372S), 5.4 µL *Bst* 2.0 DNA Polymerase (120 U/µL; New England Biolabs, M0537M), 20 µL phenol red solution (25 mM; Sigma-Aldrich, P3532), 100 µL tween 20 (20%; Sigma-Aldrich, P9416), and 331.4 µL nuclease-free water (Fisher Scientific, 43-879-36). After mixing, using a micro-pH electrode (Fisher Scientific, 11-747-328), the pH was adjusted to 7.8-7.9 using KOH (0.1 M or 1 M), to get a red solution. To prepare 200 µL LAMP master mix, we mixed 125 µL of the 2x mix, 25 µL of 10x primer mix (16 μM FIP/BIP, 2 μM F3/B3, 4 μM LF/LB) (final concentration 1.6 μM FIP/BIP, 0.2 μM F3/B3, 0.4 μM LF/LB) (Table 1), 0.67 µL *Bst* 2.0 DNA Polymerase, and 1 µL of betaine (5 M; Sigma-Aldrich, B0300-5VL), 3.13 µL bovine serum albumin (BSA) (40 mg/mL; Sigma-Aldrich, A2153), 36.0 µL trehalose (50% (w/v); Thermo Scientific Chemicals, 182550250), and 9.2 µL nuclease-free water. The pH of the master mix was adjusted to about 7.8-7.9 using KOH (0.1 M), to get a red solution. A 30 µL of the final master mix was added to each paper pad and left inside a PCR workstation (Mystaire, MY-PCR32) to dry for 2h. For each µPAD, one paper pad included a master mix with the LAMP primer mix (reaction pad) and the other one without the primer mix (no-primer control pad). When no primer mix was used, the same volume of nuclease-free water was added to the mix instead. The dried µPADs were packed in separate reclosable polypropylene bags (Uline, S-17954) and stored at −18 °C until usage. µPADs used for the field test were shipped to the field (Salinas, CA) and kept in the freezer of a household refrigerator (−18 °C) until usage.

**Table 1:**
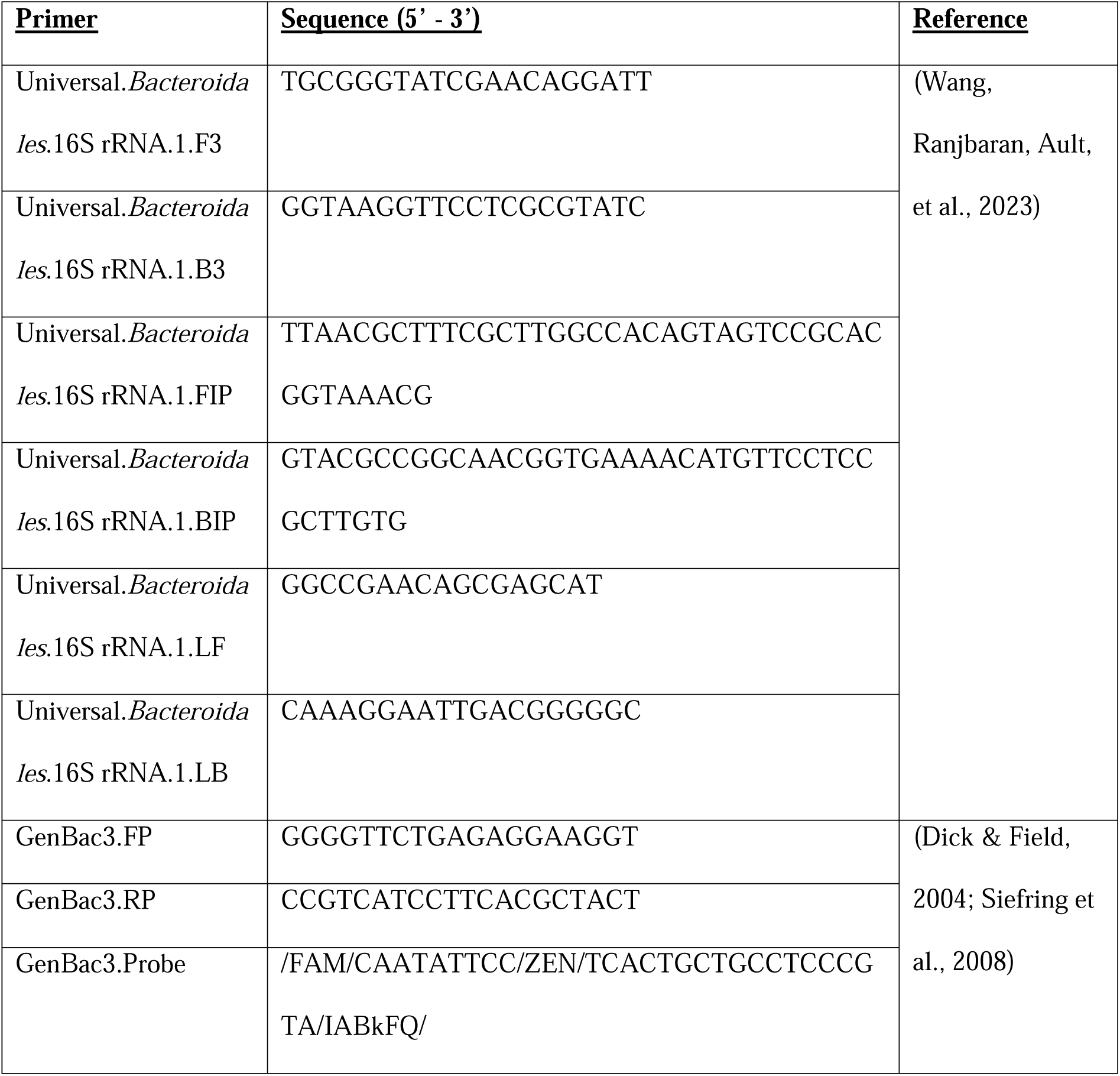
Sequences for primers and probes used in this study.

### 2.5 DNA quantification using digital PCR (dPCR)

The dPCR reactions were performed in a total volume of 40 μL, containing 10 μL 4X Probe PCR Master Mix (250102; Qiagen, USA) (final concentration 1X), 4 μL of 10X primer-probe mix (final concentration 1X, 0.8 μM forward primer, 0.8 μM reverse primer, 0.4 μM FAM probe) (Table 1), 0.5 μL EcoRI-HF restriction enzyme (NEB, R3101S), 20.5 μL nuclease-free water, and 5 μL of the template. The dPCR reactions were performed in a 26K 24-well Nanoplate (Qiagen, 250001) on a 5-plex QIAcuity One digital PCR instrument (Qiagen, 911021). The thermal cycling conditions were implemented using the following program: initial denaturation at 95 °C for 2 min, followed by 40 cycles of 95 °C for 15 s, 55 °C for 15 s, and 60 °C for 30 s. The absolute quantification method was used to calculate the DNA copy number with the QIAcuity Software Suite. The DNA sample was stored at −80 °C until usage.

### 2.6 Limit of detection (LoD) of LAMP assays using µPADs

The LAMP primer set used in this study was characterized elsewhere (Wang, Ranjbaran, Ault, et al., 2023). LoD experiments were performed on µPADs to evaluate the assay’s sensitivity (Figure 2). We used 5 levels of swine stool DNA extract concentration (50,000, 5,000, 500, 100, and 50 copies per reaction, stock quantified using dPCR) as well as no template controls (NTC) (Figure 2A). All reactions were done in triplicates. 27 µL of the template (stool DNA or nuclease-free water for NTC) was added to both paper pads (control pad and reaction pad). The µPADs were separately sealed inside resealable polypropylene bags (Uline, S-17954) and heated at 65 °C for 60 min in a water bath (Anova, ANTC01). Time-lapse video of the µPADs was taken from 0 to 60 min using a HERO8 Black digital camera (GoPro, SPJB1) (Pascual-Garrigos et al., 2021; Wang, Ranjbaran, Ault, et al., 2023). We also scanned the µPADs at 60 min using a flatbed scanner (Epson, B11B223201).

**Figure 2:**
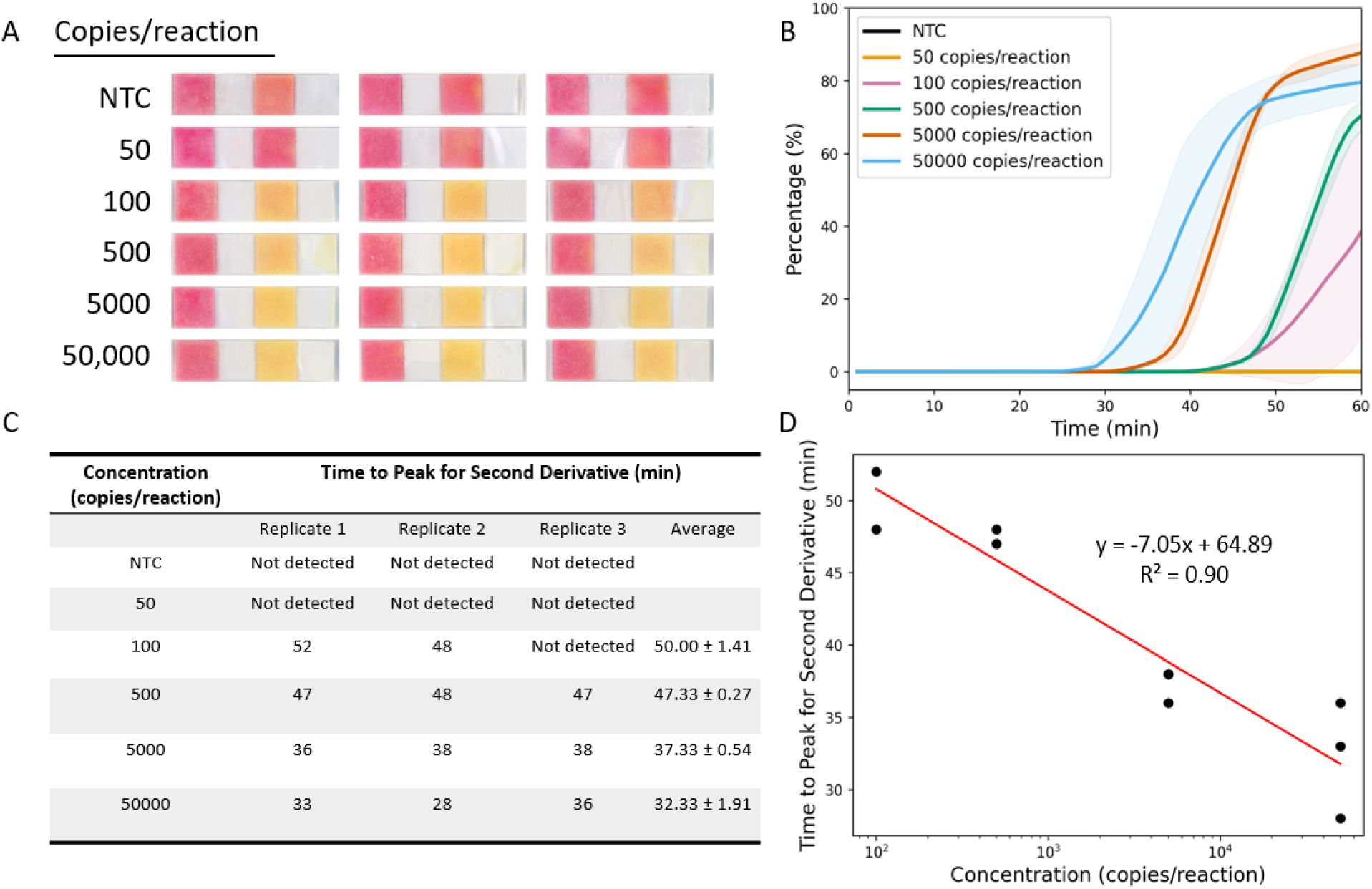
Limit of detection (LoD) and quantitative analysis of paper LAMP assays. A) Endpoint (at 60 min) result of colorimetric LoD on paper using quantified swine stool DNA at the indicated concentration (copies/reaction). The NTC replicates are reactions using nuclease-free water in lieu of DNA. B) Quantitative analysis of the LoD experiment in A using the image analysis algorithm. C) Summary table of time to peak for the second derivative calculated from B. D) Linear fit analysis generated comparing the time-to-peak for the second derivative (C) and the concentration in logarithmic scaling. A regression line was generated to quantify the concentration of the reaction using sample’s time-to-peak value.

### 2.7 Fabrication of and placement of collection flags

Following our previous results (Wang, Ranjbaran, Ault, et al., 2023), we decided to use collection flags (made from plastic sheets) to collect microbial samples in the field. These collection flags provided more consistent and reproducible LAMP results compared to lettuce leaves with grooves (Wang, Ranjbaran, Ault, et al., 2023). The collection flags were assembled using bamboo skewers (29.8 cm), transparent film (Apollo, VPP100CE), a stapler, and a paper cutter. The transparent film was pre-cut into 7.62 × 21.59 cm (3 × 8.5 inches) strips. One piece of the film was stapled together with the bamboo skewer to form a flag.

The surveyed field was labeled with row and column numbers with the distance between each row and column to be 6 meters. Samples were collected at the intersection of each row and column (approximately 100 sampling sites per acre of field) (Figure 3A). To collect airborne microbiological samples from the field, 96 collection flags were put at each sampling location seven days before sample collection. Each collection flag was encoded with a unique identifier and the location associated with the flag’s identifier was recorded.

**Figure 3:**
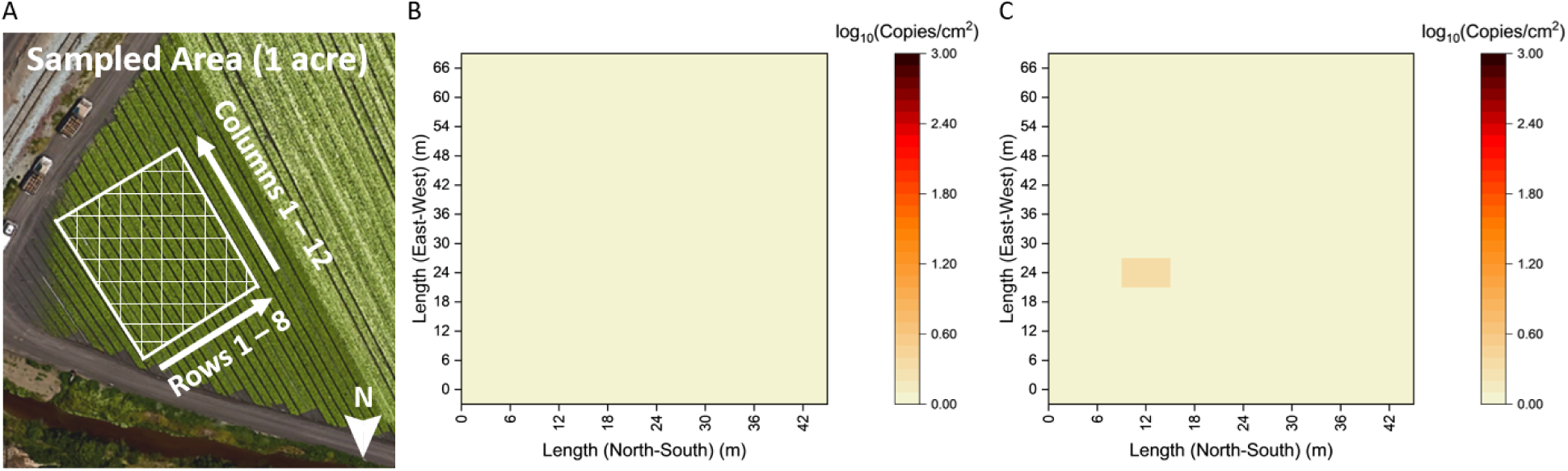
Risk of fecal contamination mapping in tested field. A) Satellite image of the sampled area. B) Risk of fecal contamination generated using paper LAMP. Quantitative analysis was performed on the paper LAMP result. The quantified target concentration was then converted into log_10_ (copies/cm^2^). C) Risk of fecal contamination mapping using qPCR The Ct value of each qPCR reaction was converted to log_10_ (copies/cm^2^) via a linear fit to log-transformed concentrations.

### 2.8 Sample collection, processing, and on-farm LAMP using the portable integrated unit

After seven days, all collection flags were collected and each flag was placed in an individual, pre-labeled Ziploc resealable storage bag (Amazon, B07NQVYCG3). Each collection flag was swabbed using a polyester-tipped swab (BD BBL, 263000) and was resuspended in 200 μL nuclease-free water (Wang, Ranjbaran, Ault, et al., 2023). To attain a more consistent swabbing, the swab was pre-wet with nuclease-free water before swabbing.

All assays were conducted in the field using FARM-LAMP. The unit was powered by a Jackery Portable Power Station (500 W, 110 V) (Jackery, Explorer 500). For each sample, the collected swab resuspension was directly transferred into a drop generator which was subsequently used to rehydrate each pad with a 27 μL drop (Ranjbaran et al., 2023). The rehydrated µPADs were sealed inside reclosable polypropylene bags (Uline, S-17954) and heated at 65°C for 60 min inside the FARM-LAMP device. The imaging system took time-lapse photos of the µPADs every 1 min during the heating time. The remaining swab resuspension samples from each location were stored in separate 1.5 mL vials, kept on ice, and shipped back to West Lafayette in a cooler box with ice packs via FedEx Priority Overnight.

### 2.9 Image Analysis Process

After the images were captured with the internal camera, they underwent a series of image processing techniques beginning with importing and resizing the first image from the test run to reduce processing time. Using Python’s OpenCV library, the GrabCut algorithm was utilized to draw the sample boundary for each paper pad and store these rectangular coordinates boundaries to create a mask for each sample. Once these masks were created for the samples in the first image, the program looped through the rest of the test images and created the mask for each sample in each image using the previous rectangular coordinates. The images of the samples were converted to the hue, saturation, value (HSV) color mode and each pixel was separated into weighted bins based on color. The color-coding function sets continuous HSV upper and lower boundaries for red, orange, and yellow based on the HSV colormap. After all pixels were identified and sorted into the different color bins ranging from dark red to light yellow, these bins were weighted based on a sigmoid function with a curve midpoint of 0.5, a curve steepness of 50, and limits of 0 to 1. The program then outputted the percentage of positivity for each sample, by calculating the ratio of the number of pixels in the weighted bins to the total number of identified pixels, to display a quantitative analysis of the color change over time (Figure 2B).

After calculating these percentages, a quantitative analysis displaying the positive percentage throughout the duration of the test run was plotted using the Matplotlib Python library. To increase the signal-to-noise ratio, a moving average with a window of 10 and a minimum period of one was applied. Then, the first and second derivatives of the positive percentage over time were calculated using Python’s Numpy gradient function. The mean y-value of the first derivative and the time to peak value for the second derivative for each concentration level and replicate were identified. The samples’ mean y-values of the first derivative were compared to the mean y-values of the lowest analyte concentration signal (eq 1 and 2) (Armbruster & Pry, 2008). If the mean y-value of the first derivative was within the lowest analyte concentration signal level, the amplification curve is then considered flat, therefore, the time to peak for the second derivative was replaced with the maximum duration of the test run, 60 minutes (Figure 2C). A calibration curve was generated comparing the time-to-peak for the second derivative and the concentration on a logarithmic scale. The linear regression equation (Figure 2D) was used to quantify the concentration of DNA for field samples.

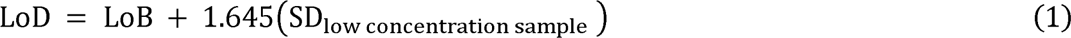

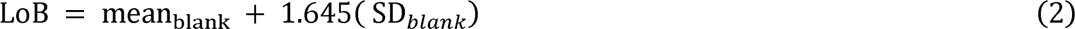

LoB: Limit of Blank; SD: Standard Deviation

### 2.10 qPCR for on-farm assay validation

qPCR assays were run on the residual swab resuspension samples from the farm, following protocols for the Luna® Universal Probe qPCR Master Mix (New England Biolabs, M3004). Each 20 µL reaction mix included 10 µL master mix, 0.8 µL PCR forward primer, 0.8 µL PCR reverse primer, 0.4 µL qPCR probe, 7 µL nuclease-free water, and 1 µL template (swab resuspension samples from the farm). The qPCR primers and probe are shown in Table 1. The assays were performed using PCR 96-plates loaded in a qTOWER³ thermocycler (Analytik-Jena, Germany) with an initial denaturation of 95 °C for 60 s and a cycling profile of 95 °C for 15 s, 55 °C for 15 s, and 60 °C for 30 s. The results of the qPCR assays were compared with those obtained from on-farm LAMP.

## 3. Results

### 3.1 FARM-LAMP fabrication and performance testing

As demonstrated in our earlier work, heating LAMP reactions under water facilitates the homogenous heat transmission required for the reactions to occur (Arumugam et al., 2020; Pascual-Garrigos et al., 2021; Wang, Ranjbaran, Ault, et al., 2023; Wong et al., 2018). Based on this knowledge, we proceeded to design and fabricate a water bath heater for conducting LAMP assays on farms. Our goal was to create a portable platform that could be used in low-resource settings, such as fields where power outlets are not available. In order to achieve this, we needed the system to have a small footprint and require a relatively low power output.

To meet these requirements, we designed FARM-LAMP to be compact with total dimensions of approximately 164 x 135 x 193 mm and a working power consumption of approximately 20W. The system consisted of a water bath, which was 3D printed with a heat-resistant resin container with a lid, a temperature control system, and an imaging system. The low power consumption allows the device to be powered with a portable power bank and the low footprint allows it to be operated easily in the back of a car in the field. Despite the low power requirement, FARM-LAMP provided fast operation, reaching 65°C in under 20 minutes. The temperature control system includes two hot rods and two water pumps to provide rapid and uniform heating. The water temperature is monitored using a waterproof digital temperature sensor and is controlled by a PID control algorithm that runs on a Raspberry Pi. The imaging system consists of a high-quality camera and four LED light sources, positioned in a way that avoids glare and reflections. To enable quantification of DNA concentration rather than just end-point determination of positivity, the camera in the imaging unit captures time-lapse images every minute, and the color change timepoint is used as input for a linear regression equation (Figure 2D) to quantify DNA concentration for unknown samples.

Overall, the FARM-LAMP device is a cost-effective and user-friendly solution for conducting LAMP assays in a range of field settings. The entire heating and imaging process is controlled by the Raspberry Pi, requiring just one button press from the user to start the process.

### 3.2 Limit of detection (LoD) and quantitative analysis of paper LAMP assays

We prepared the µPADs and dried the LAMP reaction mix on the paper strips before the experiment. Each µPAD consists of two paper pads, a control pad (does not contain LAMP primer mix) and a reaction pad (contains LAMP primer mix). The control pad serves as a negative control and is used to confirm that any positive result obtained from the reaction pad is indicative of the presence of the target and not influenced by factors such as changes in sample pH or issues related to reagent storage (Davidson et al., 2021). We prepared different concentrations of swine stool DNA and nuclease-free water for NTC. The µPADs were heated at 65 °C for 60 min in a water bath. The µPADs were scanned at 60 min using a flatbed scanner to determine the end-point result. A time-lapse video was also taken from 0 to 60 min to quantify the color-changing speed at each concentration. We determined the LoD of the µPADs as 100 copies/reaction (Figure 2).

The image analysis process was then applied to the time-lapse video. The algorithm calculated the percentage of positivity for each sample as well as the change in positivity over the course of 60 minutes. Following the calculation, a quantitative analysis displaying the positive percentage throughout the test run was plotted (Figure 2B). We calculated the second derivative of Figure 2B and adopted the time-to-peak value as the indicator for sample DNA concentration (Figure S1). In instances where the mean y-values did not surpass the lowest concentration (100 copies/reaction) signal (0.75 %positivity/min), the amplification curve is then considered flat, therefore, the sample is considered negative. Subsequently, the peak values were subjected to a linear regression analysis and the calibration curve was used to measure the concentration of DNA for field samples (Figure 2C and D).

### 3.3 Paper-based LAMP assays deployed on fresh produce farms using FARM-LAMP

The collection flags were placed in an acre of the experimental field for a period of seven days and the paper-based LAMP assay was conducted on the seventh day. All 96 collection flag samples (swab resuspension) and two positive controls (1 ng/reaction swine stool DNA extract) and two NTCs (nuclease-free water), were added on-site using the drop generator without any additional measures to avoid contamination. With a single press, the drop generator dispenses a high precision of fixed drop volume and adds it to the µPADs. The assays were conducted in the field (in the back of a car) using the portable LAMP platform powered by a Portable Power Station. Subsequently, all 100 µPADs were manually inspected after the 60-minute reaction period. We observed no color change in the NTCs, while the reaction pads of the two positive controls turned yellow. We did not observe any visible color change in either the no-primer control pads or reaction pads for the 96 collection flag samples. This indicated that all paper LAMP assays were valid and *Bacteroidales* were not detected in the fresh produce farms. The time-lapse photos of all the samples were utilized in downstream image analysis for quantitative analysis and interpretation.

As mentioned previously, the time-lapse photos of all the samples were subjected to image analysis, where the number of pixels of specific colors at each time point was quantified, and the program outputted the percentage of positivity for each sample. The positivity percentages were smoothed using a moving average filter and plotted to display a qualitative analysis of the color change (Figure S2A and C). To normalize the data, the first derivatives of the positive percentage over time were then calculated (Figure S2B and D). Additionally, the second derivative was computed to determine the exact timepoint that marked the beginning of reaction pad amplification. The mean y-values of the first derivative were compared to the mean y-values of the lowest analyte concentration signal (0.75 %positivity/min) obtained from the lowest analyte concentration (100 copies/reaction). The mean y-values for all 96 field samples and two negative controls were within the lowest analyte concentration signal level, and the amplification curve was considered flat. Therefore, the time to peak for the second derivative was replaced with the maximum duration of the test run, which was 60 minutes. The two positive controls had a mean y-value of 1.34 %positivity/min (> 0.75 %positivity/min) and were determined to be positive by the image analysis algorithm (mean positivity percentage 90.26%). These findings are consistent with our expectation that well-managed commercial fields have a low risk of fecal contamination, and therefore the concentration of *Bacteroidales* in the environment should be low (Wang, Ranjbaran, & Verma, 2023). However, trace amounts of *Bacteroidales* DNA may be present in the field due to various reasons, such as the use of organic fertilizer or residuals from previous contamination.

### 3.4 qPCR assays on the same microbial samples to validate on-farm LAMP results

The qPCR assay was performed on the same samples that were used for the on-farm LAMP assays. The qPCR assay was used to determine the ground truth concentration of *Bacteroidales* in the sample. The sensitivity of this qPCR assay was previously determined as 1 copy/reaction (Wang, Ranjbaran, Ault, et al., 2023). The comparison between LAMP and qPCR results provides an evaluation of the performance of the on-farm LAMP assay.

To generate a fecal contamination risk evaluation map, we converted the Ct value of each qPCR reaction to log_10_ (copies/cm^2^) via a linear fit to log-transformed concentrations (Wang, Ranjbaran, Ault, et al., 2023). The qPCR results showed that all 96 samples had a *Bacteroidales* concentration of 0 – 2.24 copies/cm^2^ (Figure 3C). This result is consistent with the LAMP assay results, which showed < 3 copies/cm^2^ (or <100 copies/reaction). The complete agreement between the two assays indicates that the on-farm LAMP assay has reasonable performance for *Bacteroidales* detection in fresh produce farms. Although the *Bacteroidales* LAMP assay is not as sensitive as qPCR, it is still sufficient for identifying possible fecal contamination events in the field. Furthermore, the on-farm LAMP assay offers the advantages of being rapid, low-cost, and easy to use, which makes it a promising tool for microbial detection in field settings.

## 4. Discussion

### 4.1 Needs and challenges of bringing nucleic-acid testing to fresh produce farms

A number of commercial portable instruments are available to perform food pathogens nucleic-acid testing for the fresh produce industry. Examples of these devices include the 3M Molecular Detection System (3M, USA) to detect *Salmonella spp.* and *Listeria monocytogenes*, HumaLoop T and HumaLoop M (HUMAN Diagnostics, Germany) for the detection of *Mycobacterium tuberculosis,* and *Plasmodium spp*., and Genie II (OptiGene, UK) for detection of plant pathogens. However, many of these assays are complex to operate and necessitate specialized personnel for reagent handling and sample preparation, making them unsuitable for use in the field. Consequently, food microbial testing still requires sending samples to centralized labs for analysis. Not only do courier fees for sample shipping contribute significantly to the cost of microbiology testing services, but the lab-based services also result in extended turnaround times for test results ranging from 1 to 3 days. This delay can significantly impact the decision-making process for marketing the tested batch of product, which is especially crucial for fresh produce as it has a limited shelf life.

*Bacteroidales* as a FIB has been extensively used to evaluate fecal contamination both in fresh produce fields and for water quality monitoring (Jiang et al., 2018; Mascorro et al., 2018; Ravaliya et al., 2014; Wang, Ranjbaran, Ault, et al., 2023). In the previous studies, we have determined that there is a ∼10^4^-fold difference in *Bacteroidales* concentration between fields with a high risk of fecal contamination (adjacent to animal units) (Wang, Ranjbaran, Ault, et al., 2023) and a low risk of fecal contamination (commercial fresh produce fields) (Wang, Ranjbaran, & Verma, 2023). The significant difference in *Bacteroidales* concentrations between “high risk” and “low risk” fields makes it a valuable tool for researchers and industry stakeholders to monitor fecal pollution and facilitate microbial safety in the fresh produce industry. If a portable and easy-to-use *Bacteroidales* assay is available, it could be adopted by industrial stakeholders as a complementary tool for pre-season and pre-harvest decision-making in fresh produce operations. During the pre-season period, these tools could be used to assess the risk of fecal contamination from nearby animal operations. During the pre-harvest period, this tool can assist in determining whether the product in this site is safe to harvest and send to market.

### 4.2 Enabling paper-based LAMP assays in the farm using the current integrated unit

In this study, we have focused on enabling an easy-to-use paper-based biosensor with a portable testing unit for assessing fecal contamination on farms. We developed a fully-integrated LAMP testing platform – FARM-LAMP, encompassing heating, imaging, fluid delivery, and paper-based LAMP assay, which we subsequently deployed on a commercial lettuce farm.

We successfully integrated the developed *Bacteroidales* LAMP assay into µPADs, which were comprised of pre-fabricated paper strips featuring one no-primer control pad and one reaction pad (Davidson et al., 2021; Wang et al., 2021; Wang, Ranjbaran, Ault, et al., 2023). All necessary reagents were pre-dried onto the paper strips, thereby enabling end-users to directly apply their samples onto the respective pads, and initiate the reaction. As such, our paper-based LAMP method streamlines the testing process, reduces the number of handling steps required, and enhances the usability of nucleic acid tests for non-specialists operating in the field. To enhance the overall user experience of the assay, we designed and fabricated a drop generator that enables precise sample volume delivery to the paper-based biosensor via a simple button press (Ranjbaran et al., 2023). This feature obviates the need for costly commercial micro-pipettors, thus increasing the accessibility and affordability of the assay. Furthermore, the disposable nature of the drop generator effectively mitigates the risk of cross-contamination during use by non-specialists.

We engineered a compact and portable heating and imaging unit to facilitate uniform heat distribution for the LAMP assay. Our heating component exhibited rapid and efficient operation, capable of reaching the programmed temperature of 65°C within 20 minutes and maintaining this temperature throughout the course of the test. Concurrently, the imaging unit integrated within the device captured time-lapse images every minute to monitor and record any color changes occurring during the assay. Following image capture via the camera, a series of image processing techniques were employed to ascertain the sample’s result, either positive or negative, and its concentration of *Bacteroidales* if deemed positive. This analysis was accomplished via quantitative assessment of the color change of each pad and computation of the percentage of positivity for each sample over time. The positivity threshold time point was then correlated with a predefined calibration curve to obtain an accurate quantitative concentration result. By adopting this approach, we not only provided end-users with more comprehensive assay results but also mitigated potential color perception discrepancies among individuals. Although the current image analysis is performed on a separate computer, we anticipate integrating the entire process onto the device’s Raspberry Pi for real-time execution in the future. By transitioning to this approach, we aim to further expedite the analysis process and enhance the overall efficiency and user-friendliness of the assay.

Our integrated platform successfully generated quantitative results within an hour of sample collection. We deployed the unit on a commercial lettuce farm, operating from the back of a vehicle and powered by a portable power station, to detect *Bacteroidales* as a biomarker of fecal contamination. The assay achieved a sensitivity of approximately 3 copies of *Bacteroidales* per cm^2^ of the collection flags’ surface (corresponding to 100 copies/reaction). In comparison to the liquid LAMP assay (LoD 50 copies/reaction), the paper-based LAMP exhibits a slightly worse LoD (Wang, Ranjbaran, Ault, et al., 2023). However, due to a higher sample volume was added to the paper LAMP assay (adding 27 µL for paper LAMP and 1 µL for liquid LAMP), its final LoD in copies of *Bacteroidales* per cm^2^ of the collection flags’ surface is marginally better than the liquid LAMP assay (13.5× more sensitive). In addition, the multiple liquid handling steps and low volume pipetting (0.5 – 2.5 µL) necessary for the liquid LAMP assay render it impractical for non-specialist users. In contrast, the paper-based LAMP assay, with comparable sensitivity, represents a more viable option for on-field testing scenarios.

To our knowledge, this investigation represents the first demonstration of a portable paper-based LAMP testing platform implemented on a fresh produce farm. This development holds significant promise for incorporating future NAATs as part of standard growing and harvesting practices within the fresh produce industry.

## 5. Conclusions

Here, we have developed a fully integrated paper-based LAMP testing platform that can be used on-site for the detection of *Bacteroidales* as a fecal contamination biomarker. Our platform offers the following five advantages: i) it can provide quantitative results within an hour of sample collection; □) it performs similarly to a qPCR assay in terms of analytical sensitivity and specificity with non-enriched samples; □) it is highly versatile, with the ability to operate in the back of a vehicle; □) it is user-friendly and could be used by non-specialist users; □) it provides end-users with quantitative results (e.g., concentration of target) and mitigates potential color perception discrepancies among individuals.

The current approach has two limitations: i) the sample must be processed and added to each reaction zone separately (the development and incorporation of a fluid separation system would allow the user to add the sample to a single site and the sample would be distributed evenly to all reaction zones); ii) the current image analysis is performed separately after the assay (the entire process needs to be incorporated on the FARM-LAMP device for real-time execution).

As a demonstration, we successfully deployed the unit on a commercial lettuce farm and generated a heatmap to showcase the distribution of fecal contamination within a 1-acre field. The integrated platform showed a concordance of 100% when compared to the standard lab-based tests. This could be adopted by industrial stakeholders as a complementary tool for decision-making during pre-seasons and pre-harvests of fresh produce. Additionally, it allows for a better understanding of the extent of contamination in the fields, which can be used to identify problematic areas and develop appropriate measures for remediation.

In the majority of cases, enteric pathogens exhibit a notably low concentration and are known to be distributed heterogeneously within food products (Jongenburger et al., 2011). However, regulatory requirements for the detection of these pathogens are exceedingly stringent, mandating the identification of a single live bacterium in a 25-gram sample. Consequently, to reduce false-negative rates, direct pathogen detection needs to increase the sampling number within a batch of food (increasing cost) or enhance the sensitivity of the detection assay. Unfortunately, no currently available technology can achieve the required high sensitivity of 4.4×10^-6^ copies/µL in a timely and cost-effective manner (Alafeef et al., 2020; Jasim et al., 2019; Shin et al., 2018; Trinh et al., 2019; Wu et al., 2022). Consequently, the adoption of such assays and technologies by the industry remains unfeasible.

From the perspective of the authors, relying solely on one testing approach to meet the required sensitivity for detecting foodborne pathogens appears impractical. Therefore, direct pathogen screening may not be the most effective method, and instead, the use of indicators seems to be a more feasible option to assess the potential risk of contamination. In addition, it is difficult to determine the presence/absence of all potential pathogens that are present, and the indicators quantification approach simplifies the testing process by only quantifying one group of microorganisms and providing risk evaluation of the presence of pathogens. Among the available indicators, *Bacteroidales* are particularly promising due to their high concentrations in feces (total concentration 10^11^ bacteria/g of stool) (Gorbach, 1996), accounting for about 30%–40% of total fecal bacteria (Mascorro et al., 2018), which is several orders of magnitude higher than the concentration of pathogens, making them easier to detect and more homogeneously distributed in the event of fecal contamination. However, it is crucial to acknowledge that the indicator approach has its limitations. While the quantification of *Bacteroidales* provides valuable risk assessment, it does not directly correlate with the presence of specific pathogens in the sample. Instead, it offers a risk-based evaluation. As a result, end users should utilize the obtained results to implement appropriate safety practices and protocols to mitigate potential sources of contamination based on the assessed risk.

Overall, this investigation is the first to demonstrate a portable paper-based LAMP testing platform implemented on a fresh produce farm. Our integrated paper-based LAMP testing platform holds great promise as a reliable and convenient tool for on-site NAATs. We anticipate that this advancement will promote the adoption of NAATs as part of standard growing and harvesting practices within the fresh produce industry. We also hope that this work can stimulate further research in the development of on-farm diagnostic tools for other agricultural applications.

## Acknowledgment

This work was funded in part by the Center for Produce Safety (CPS Award Number: 2021CPS12), the California Department of Food and Agriculture (CDFA Agreement No. 20-0001-054-SF), and the U.S. Department of Agriculture’s (USDA) Agricultural Marketing Service (USDA Cooperative Agreement No. USDA-AMS-TM-SCBGP-G-20-0003). The project entitled Field evaluation of microfluidic paper-based analytical devices for microbial source tracking was funded in whole or in part through a subrecipient grant awarded The Center for Produce Safety through the California Department of Food and Agriculture 2020 Specialty Crop Block Grant Program and the U.S. Department of Agriculture’s (USDA) Agricultural Marketing Service. We acknowledge support from The Center for Produce Safety (2023CPS11) and the U.S. Department of Agriculture’s (USDA) Agricultural Marketing Service through grant AM22SCBPCA1133 for “Testbeds for microbial source tracking using microfluidic paper-based analytical devices”. Any opinions, findings, conclusions, or recommendations expressed in this publication or audiovisual are those of the author(s) and do not necessarily reflect the views of The Center for Produce Safety, the California Department of Food and Agriculture, or the U.S. Department of Agriculture (USDA).

We are grateful for the collaboration with Patricia Contreras, Felice Arboisiere, and Natalie Dyenson from Dole Fresh Vegetables, Inc. and Dole Food Company, Inc. for their assistance in collecting samples in the field.

The plasma treatments were performed in Prof. Rahim Rahimi’s laboratory located at the Birck Nanotechnology Center of Purdue University. Sina Nejati, his graduate student, provided initial training to use the plasma generator.

