## Supporting Information for "A portable, easy-to-use paper-based biosensor for rapid in-field detection of fecal contamination on fresh produce farms"

**for**

**Supporting Figure**

Figure S1: Second derivative of the limit of detection experiment. The second derivative (red line) of the positive percentage (black line) throughout the test run is plotted against time. The time-to-peak value for the second derivative was adopted as the indicator for sample DNA concentration.

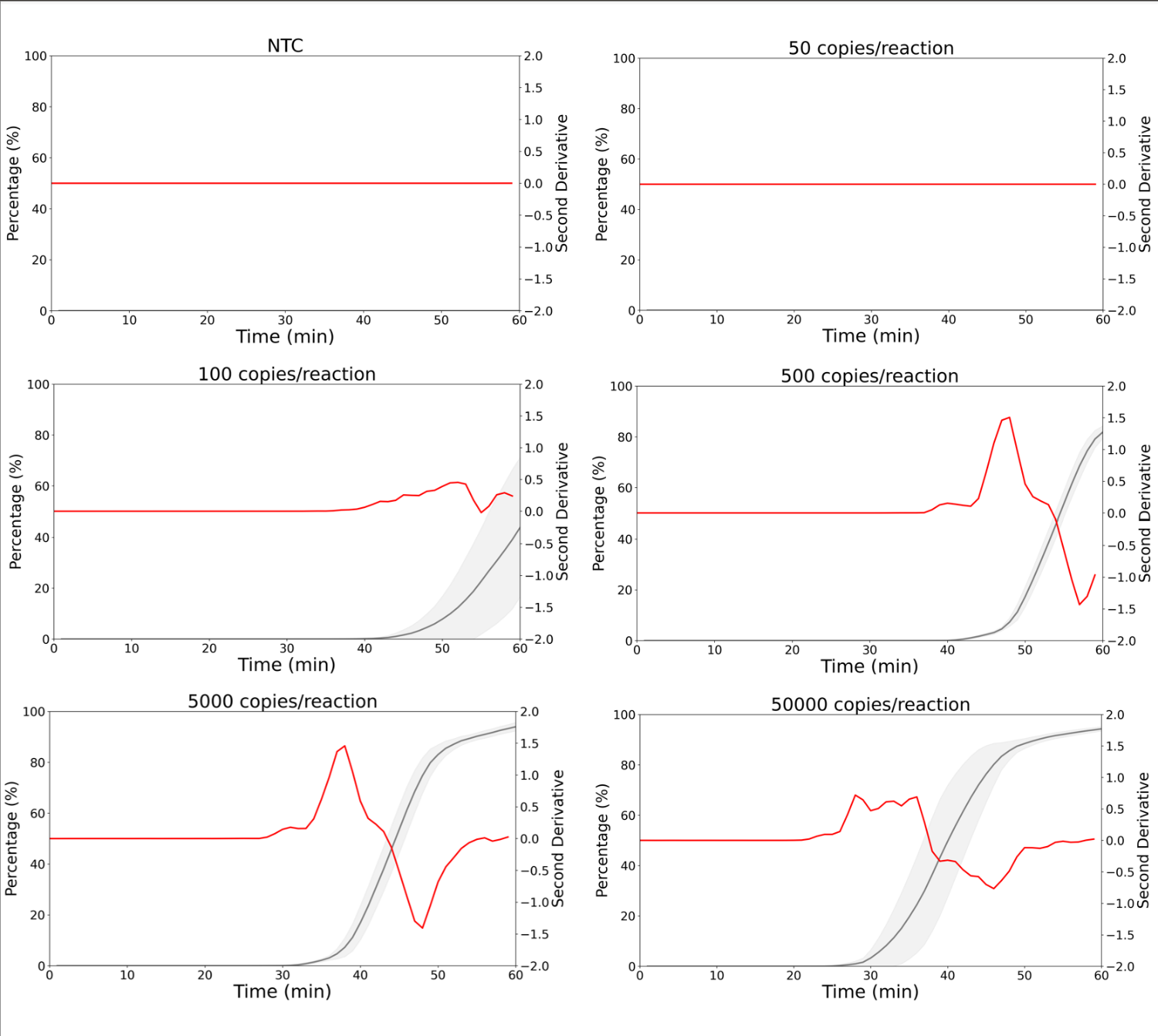

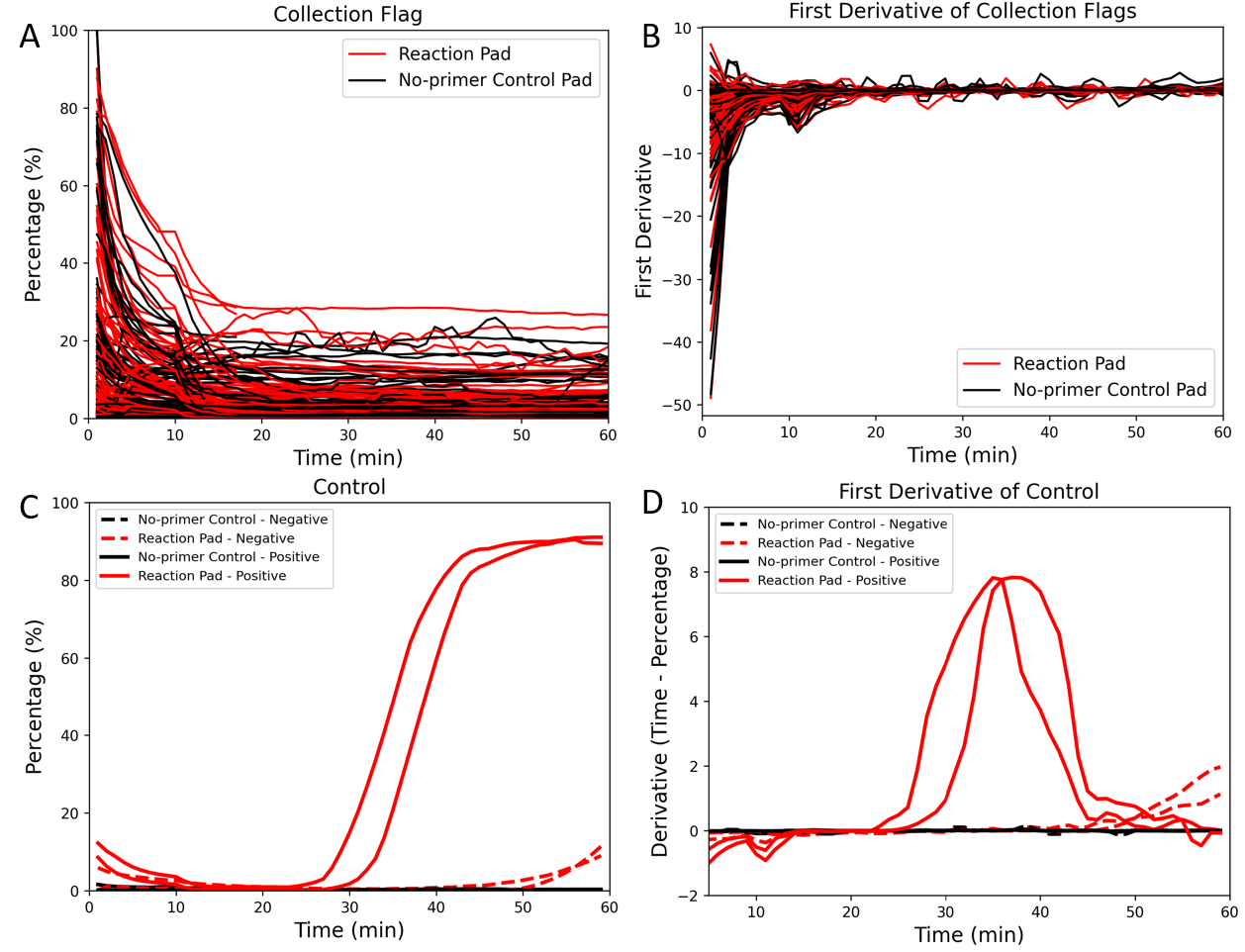

Figure S2: Image analysis result of the field experiment. A) The positivity percentage result overtime for each field sample. The curves were smoothed using a moving average filter and plotted to display a qualitative analysis of the color change. B) First derivative of the positive percentage in A. C) The positivity percentage result overtime for each control sample. The curves were smoothed using a moving average filter and plotted to display a qualitative analysis of the color change. D) First derivative of the positive percentage in C.
